# Assessment of Functional Connectome Construction Strategies in Neurodegeneration

**DOI:** 10.1101/385385

**Authors:** J Vanhoecke, P McColgan, A Razi, S Gregory, K Seunarine, A Durr, R Roos, B Leavitt, RI Scahill, C Clark, SJ Tabrizi, G Rees

**Affiliations:** Department of Neurodegenerative Disease, UCL Institute of Neurology, London, WC1N 3BG, UK; Wellcome Trust Centre for Neuroimaging, UCL Institute of Neurology, London, WC1N 3BG, UK; Department of Electronic Engineering, NED University of Engineering and Technology, Karachi, Pakistan; Developmental Imaging and Biophysics Section, UCL Institute of Child Health, London, WC1N 1EH, UK; Institut du Cerveau et de la Moelle épinière (ICM), Genetic department, AP-HP, Sorbonne Université, Inserm U 1127, CNRS UMR 7225, Pitié-Salpêtrière University Hospital, Paris, France; Department of Neurology, Leiden University Medical Centre, 2300RC Leiden, The Netherlands; National Hospital for Neurology and Neurosurgery, Queen Square, London, WC1N 3BG, UK

**Keywords:** Functional connectomics, modularity, machine learning, Huntington’s disease, graph theory

## Abstract

Connectomics can be used to investigate functional brain networks in neurodegenerative diseases including Huntington’s disease (HD). In this developing field, different connectome construction strategies have emerged in parallel. However, there is a need to understand the influences of different strategies on subsequent analyses when constructing a connectome. This study systematically compares connectome construction strategies based on their biological relevance to functional networks in neurodegeneration.

We asked which functional connectome construction strategy was best able to discriminate HD gene carriers from healthy controls, and how such a strategy affected modular organization of the network. The major factors compared were principal component-based correction versus wavelet decomposition for physiological noise correction, the type of parcellation atlas (functional, structural and multi-modal), weighted versus binarized networks, and unthresholded versus proportionally thresholded networks. We found that principal component-based correction generated the most discriminatory connectomes, while binarization and proportional thresholding did not increase discrimination between HD gene carriers and healthy controls. When a functional parcellation atlas was used, the highest discrimination rates were obtained. We observed that the group differences in modular organization of the functional connectome were greatly affected by binarization and thresholding, showing no consistent pattern of modularity.

This study suggests that functional connectome construction strategies using principal component-based correction and weighted unthresholded connectivity matrices may outperform other strategies.

## Introduction

Functional and structural brain networks can be constructed using resting state fMRI and diffusion tractography respectively. For functional connectomics, different methodologies have been used to construct and compare brain networks (Sporns, 2011). Amongst these factors some of the important choices are the type of physiological noise correction (Marchitelli *et al.*, 2016), the size and nature of the parcellation atlas (Arslan *et al.*, 2017), and the application of thresholding and binarization of connectivity matrices (Garrison *et al.*, 2015). However, comparative studies of the interactions of these factors when constructing the connectome have not been performed. Understanding the effects of connectome construction strategies is important to provide insights when developing a standard for the field.

One difficulty in evaluating connectome construction strategies is the lack of a gold standard. To overcome this, we set out to investigate how different strategies affect the ability of using a constructed connectome to distinguish between Huntington’s disease (HD) gene carriers and healthy controls. HD is an autosomal dominant neurodegenerative disease, and is fully penetrant in those with greater than 39 cytosine-adenine-guanine (CAG) repeat expansions in the HD gene on chromosome 4, see McColgan 2018 for a review (McColgan and Tabrizi, 2018). Thus HD gene testing allows us to identify with certainty those who will develop the disease. In this context, genetic testing can be used as a diagnostic gold standard. This then allowed us to evaluate connectome construction strategies in their ability to discriminate between HD and healthy controls.

Options for physiological noise correction include band pass filtering and regression of white matter, CSF and global signal. More recently, due to concerns regarding global signal regression (Carbonell, Bellec and Shmuel, 2014) more conservative approaches use only the principal components of white matter and CSF signals. This approach was developed as the anatomical CompCor method (Behzadi *et al.*, 2007) and is implemented in the freely available Conn functional connectivity toolbox (Whitfield-Gabrieli and Nieto-Castanon, 2012). Wavelet decomposition can also remove signal attributable to physiological noise. Here, the signal is broken down into its underlying constituent frequencies by scaling and shifting of a brief oscillation, a so-called wavelet. By applying the maximal overlap discrete wavelet transform (MODWT) with Daubechies wavelets (Daubechies, 1988) to the raw rs-fMRI time series, correlation matrices are formed. Brainwaver is a freely available R-based package that can be used to perform such a wavelet decomposition (Achard *et al.*, 2006).

Following physiological noise correction, the brain is parcellated into discrete regions, such that each brain region acts as a node in the network and the temporal correlations of the fMRI time series between regions act as functional connections or edges. Atlases can be defined anatomically, such as the Desikan-Killiany atlas where labeling is inferred from anatomic curvature (Desikan *et al.*, 2006). Functional atlases can also be defined based on resting state connectivity (Yeo *et al.*, 2011). More recently multimodal atlases have been developed, such as the Glasser atlas (Glasser *et al.*, 2016), which is based on task fMRI, rs-fMRI, and cytoarchitectonic features. However, currently there is no consensus in the literature as to the optimal strategy for brain parcellation (Arslan *et al.*, 2017).

Following the parcellation of the brain into various nodes or brain regions, thresholding is often performed in order to remove spurious connections. Binarization can also be carried out, such that the connection is either present or absent with the magnitude of the temporal correlation ignored. As there is no consensus about what threshold value to use, often a range of different thresholds are applied (Bullmore and Sporns, 2009). A variety of different threshold approaches have been described in the literature including absolute, proportional, consistency and consensus thresholding, but no optimal method has been identified (Qi *et al.*, 2015). When thresholding is applied, it is often combined with a minimum spanning tree to retrieve the backbone of the graph (Alexander-Bloch *et al.*, 2010), which is a subgraph with minimal connection weights, while ensuring all nodes are connected to the network. Alternatively, thresholding can be avoided by using weighted connectivity matrices, using the raw correlations (Bullmore and Sporns, 2009).

Machine learning tools such as support vector machines can determine a discrimination rate between two groups, as implemented in for example the Pattern Recognition for Neuroimaging Toolbox, PRoNTo (Schrouff *et al.*, 2013). This has been used previously to classify structural networks from different Brain-derived Neurotrophic Factor (BDNF) genotypes from healthy subjects (Ziegler *et al.*, 2013).

Modularity is a measure of functional segregation and represents how well the network can be divided into distinct cooperating units, also called modules or communities (Rubinov and Sporns, 2010). In neurodegeneration, changes occur in the modular organization of the brain (McColgan *et al.*, 2015) and therefore this is a complementary way to assess the effect of pathology on the brain network.

Here we set out to address two specific questions: first, which functional connectome construction is most able to discriminate HD gene carriers from healthy controls? Second, how does the method of functional connectome construction affect determination of modular organization? For both research questions, we undertook a systematic comparison considering different factors and their interactions. These factors were principal component-based correction versus wavelet decomposition for physiological noise correction, the type of parcellation atlas (functional, structural and multi-modal), weighted versus binarized networks, and unthresholded versus proportional thresholded networks, resulting in a systematic comparison of 66 connectome construction strategies (Figure 1).

**Figure 1:**
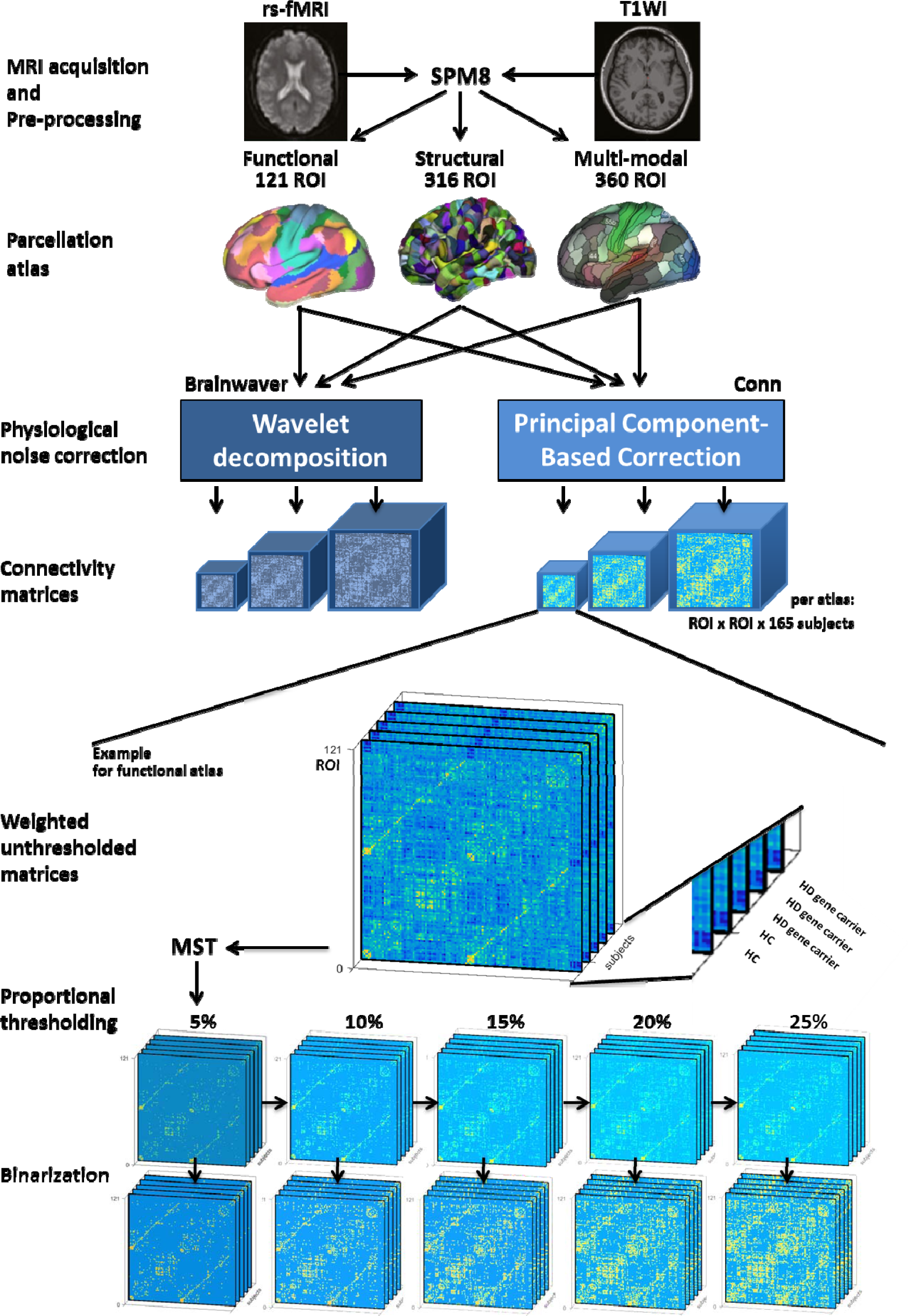
Overview systematic comparison of connectome construction strategies. Rs-fMRI was obtained from 86 HD gene carriers and 79 healthy controls. Functional images were realigned and then coregistered to anatomical images. ROIs of three parcellation atlases were coregistered to the new anatomical images. For the use of each atlas a wavelet decomposition or component-based correction was applied for physiological noise correction. This resulted in weighted unthresholded matrices, whereof the minimum spanning tree (MST) is retrieved and subsequently, the strongest connections are added back to obtain a range of proportionally thresholded matrices. Finally, each of these thresholded matrices have been binarized. The following pictures are adapted from original publications: MRI acquisition (McColgan, Gregory, et al. 2017); functional (Yeo et al. 2011), structural (Romero-Garcia et al. 2012) and multi-modal (Glasser et al. 2016) parcellation atlas. HC: healthy controls. HD: Huntington’s disease. MST: minimum spanning tree algorithm. ROI: region of interest. rs-fMRI: resting-state functional Magnetic Resonance Imaging. SPM8: statistical parametric mapping. T1WI: T1-weighted imaging.

To our knowledge, there is no systematic comparison of the effect of all factors mentioned above on functional brain networks, especially not applied in a clinical population. While optimization of connectome construction protocol has been done before, it mainly focused on one single factor, for instance the effect of thresholding, ignoring other influences (Garrison *et al.*, 2015). As these factors could have critical interactions in determining an optimal strategy, a systematic comparison can be particularly valuable.

## Materials and Methods

### Cohort

We assembled a cohort that comprised HD gene carriers and healthy controls from the final visit (2014) of the prospective Track-On HD study (Klöppel *et al.*, 2015). From a total of 243 eligible participants, 78 participants were excluded due to poor quality rs-fMRI data (Klöppel *et al.*, 2015). If not further specified, the remaining cohort of 165 participants was used in full. For the classification analysis, a subdivision was taken, so that the number of participants in each group was equal. Participants were pairwise matched for age and gender, so that each match consisted of one HD gene carrier and one healthy control participant from the same gender and with an age difference spanning less than 1 year. This strategy yielded 49 HD gene carriers and 49 HC (Table 1), without significant differences in age (2 tail t-test, p=0.93), gender (*a priori* equal), education (WMW, p=0.38) and study site (Chi-square, p=0.91). This subdivision is further referred to as a pairwise matched cohort. An alternative and less stringent cohort subdivision was also performed, in which cohorts were selected by median age, see Supplementary Methods.

**Table 1:**
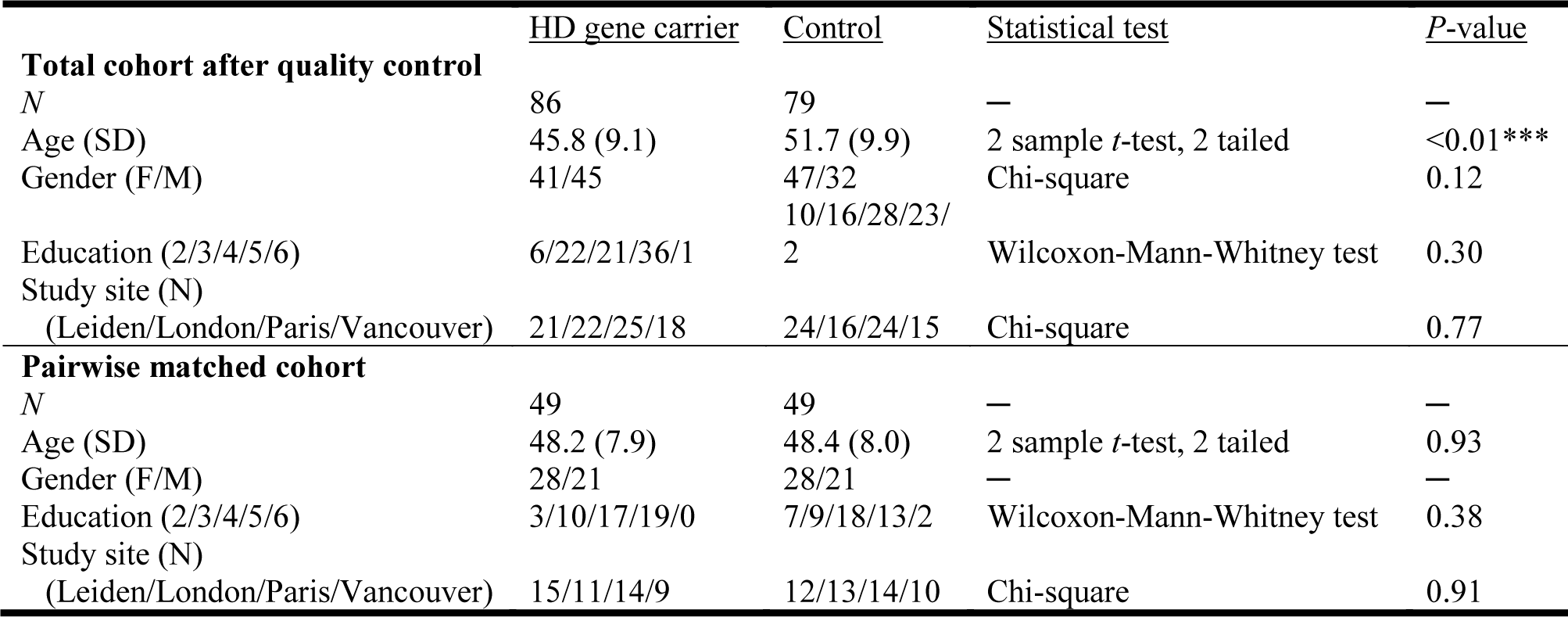
**Cohort demographics.** Values are mean (standard deviation) or n/n. See text for composition strategy.

### MRI acquisition

T1-weighted and rs-fMRI data were acquired on two different 3T MRI scanner systems (Philips Achieva at Leiden and Vancouver and Siemens TIM Trio at London and Paris). Scanning time was approximately 12 min for the T1-weighted acquisitions and 8 min for the rs-fMRI. For the rs-fMRI T2*-weighted echo planar imaging sequence with a repetition time of 3 s was used, resulting in 165 whole-brain time series. For both Siemens and Philips scans the voxel size was 3.3 × 3.3 × 3.3 mm^3^. Detailed information on parameters and control procedures are provided in the Track-On HD study (Klöppel *et al.*, 2015; McColgan *et al.*, 2017).

### Data Pre-processing

Pre-processing of the structural and functional MRI data was performed with MATLAB R2016a, making use of SPM8 (Friston *et al.*, 2011) and Conn (Whitfield-Gabrieli and Nieto-Castanon, 2012). Each stage of the pre-processing was subjected to the quality control procedures of Track-On HD (Klöppel *et al.*, 2015). T1-weighted images were segmented using three different parcellation atlases: a combined functional atlas containing cortical (Yeo *et al.*, 2011) and sub-cortical regions (Choi, Yeo and Buckner, 2012), a structural atlas (Romero-Garcia *et al.*, 2012) and a multi-modal atlas (Glasser *et al.*, 2016). Atlases were registered to the anatomical image using NiftiReg (Modat *et al.*, 2010). For the Romero atlas the globus pallidus, amygdala, nucleus accumbens and cerebellum were excluded. The functional, structural and multi-modal parcellation strategies entailed 121 ROIs, 316 ROIs and 360 ROIs respectively.

Functional images were realigned, then corrected for estimated head movement, and coregistered to the new anatomical image. For each participant, a regional mean time series of the resting-state were obtained by averaging the fMRI time series over each voxel of a ROI. Physiological noise correction was then performed as detailed below.

### Physiological noise correction

To perform a wavelet decomposition, six motion parameters were first linearly regressed out from the time series (Achard *et al.*, 2006). Residuals were decomposed with Brainwaver (http://cran.r-project.org/web/packages/brainwaver/index.html) in RStudio 0.98.1078, using Daubechies wavelets, and the correlation matrix generated in the 4^th^ wavelet scale (0.03-0.06 Hz) was retrieved for further analysis, in keeping with the literature (Achard *et al.*, 2006). The use of higher frequencies from lower wavelet scales has been reported (Richiardi *et al.*, 2011; Vértes *et al.*, 2016). However, frequencies of 0.1-0.5 Hz are thought to originate from respiration (Cordes *et al.*, 2001) and thus, to make an unbiased comparison with bandpass-filtering used in CompCor, the lowest wavelet scale corresponding to a frequency below 0.1 Hz was used.

An alternative physiological noise correction was performed using Conn (http://www.conn-toolbox.org). Using a CompCor method (Behzadi *et al.*, 2007), regression of the first 5 principal components of the white matter and cerebrospinal fluid (CSF) signal was performed along with 6 movement parameters (Whitfield-Gabrieli and Nieto-Castanon, 2012). Global signal regression was not performed. Subsequently, a bandpass filter of 0.01-0.10 Hz was applied. Bivariate correlations were retrieved after a Fisher transform was performed to improve the normality assumption of the distribution of the correlation coefficients (Whitfield-Gabrieli and Nieto-Castanon, 2012).

### Connectome construction strategies

First, a minimum spanning tree (MST) algorithm was applied to the raw weighted and unthresholded connectivity matrices (CM) to retrieve a maximally sparse matrix while ensuring all nodes are connected to the network. This means that graph subset is the matrix backbone that has a minimum possible edge weight and does not include cycles. Subsequently, connections were added back to the network in decreasing strength to obtain weighted proportionally thresholded CM with a density of 5%, 10%, 15%, 20% and 25%. The second series of 5 CM was obtained by binarization of the weighted and proportionally thresholded CM (Rubinov and Sporns, 2010). This resulted in 1 weighted and unthresholded; 5 weighted and proportionally thresholded; and 5 thresholded and binarized CM in total. Because these 11 CM were used for each of three parcellation atlases and two physiological noise correction methods resulting in 66 connectomes per participant in total (see also Figure 1).

### Classification with support vector machine

For each of the connectome as described above, the connectomes of pairwise matched HD gene carriers and healthy controls were classified using Pattern Recognition for Neuroimaging Toolbox v2.01 (PRoNTo), http://www.mlnl.cs.ucl.ac.uk/pronto (Schrouff *et al.*, 2013). Leave-one-out cross-validation (100 permutations) was applied. A post-hoc analysis was performed to assess effects of overall functional connectivity and motion (see Supplementary Methods).

### Modularity

For each connectome construction strategy, the community affiliation vector was computed for the group averaged CM of HD gene carriers and for the group averaged CM of HC, using the Louvain algorithm for community detection (Blondel *et al.*, 2008). 250 iterations were performed with a resolution parameter γ = 1 (default), followed by a consensus clustering (Lancichinetti and Fortunato, 2012). The modules for brain networks were visualized with BrainNet Viewer (Xia, Wang and He, 2013). Two related measures of modularity are the participation coefficient (PC); a measure that quantifies how evenly the node’s connections are distributed across modules (Sporns and Betzel, 2016) and within module Z-score; a way to express intra-modular connectivity (Fornito, Zalesky and Breakspear, 2015). The rationale for using these measures was that it allows a quantitative comparison of the hubness of networks in addition to a qualitative description of a network using modularity, which are both important in functional imaging (Power and Schlaggar, 2013). PC and within module Z-score were computed as implemented in the Brain Connectivity Toolbox version 2017-01 (https://sites.google.com/site/bctnet/). For these measures, differences between HD gene carriers and HC were assessed using permutation testing (10 000 iterations). Age, gender, site of acquisition, education and overall functional connectivity (FC) were regressed out as covariates and false discovery rate (FDR) correction was applied. The nodes that were identified as significant (p-value<0.05, FDR corrected) for the PC and within module Z-score separately were contrasted in both the number as location of these nodes across the different connectome construction strategies. Nodes were visualized with BrainNet Viewer (Xia, Wang and He, 2013).

## Results

### Classification of HD gene carriers and healthy controls

The first aim of this study was to assess which functional connectome construction method was most discriminative for a machine learning classification of HD gene carriers and healthy controls (HC). Regardless of thresholding or binarization, the classification for the pairwise matched cohort yielded a higher area under the curve (AUC) for physiological noise correction performed using Conn connectomes, ranging from 0.58 to 0.78, compared to Brainwaver connectomes, ranging from 0.32 to 0.57 (Figure 2). The AUC using the functional, multi-modal or structural parcellation atlas were 0.78, 0.70 and 0.65 respectively. For weighted matrices the functional atlas consistently gave for each threshold a higher AUC compared to the other atlases. The AUC for the functional atlas using Conn ranged from 0.69 to 0.78, compared to a range of 0.60 to 0.70 for the multi-modal and 0.59 to 0.65 for the structural atlas. For binarized matrices using Conn the AUC for the functional atlas ranged from 0.64 to 0.71, while the AUC ranges for the multi-modal and the structural atlas were 0.58 to 0.68 and 0.58 to 0.63 respectively. Therefore, thresholding or binarization had a minimal effect on the AUC.

**Figure 2:**
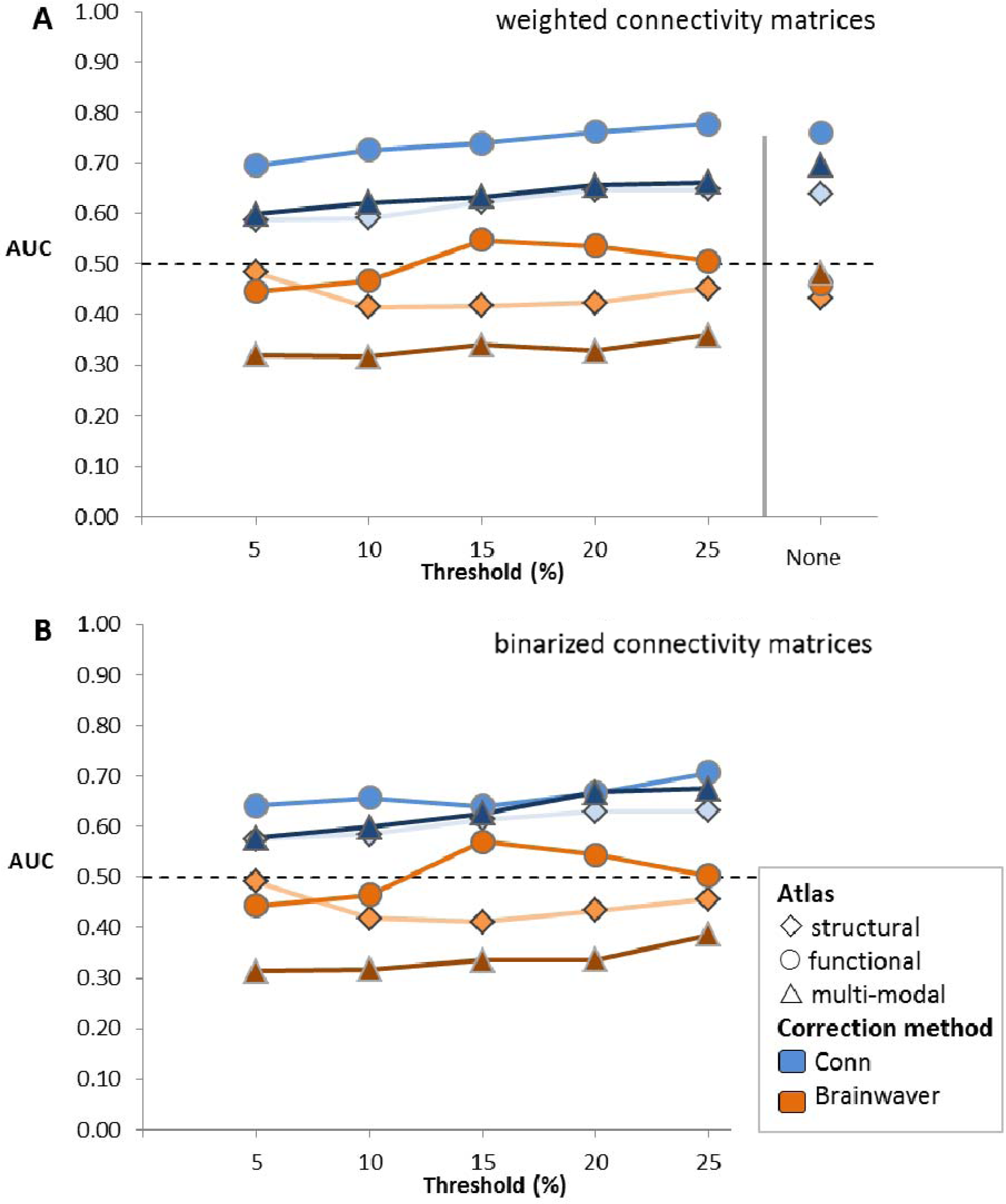
AUC for pairwise matched cohort. The connectivity matrices of 49 HD gene carriers and 49 healthy controls pairwise, matched for gender and age, are classified with PRoNTo in a leave-one-out cross-validation. The area under the curve (AUC) is obtained for each of 66 connectome construction strategies. The structural atlas is based on anatomical demarcations (Romero-Garcia et al. 2012), the functional atlas is based on intrinsic connectivity of brain regions (Yeo et al. 2011), and the multi-modal atlas made use of both criteria (Glasser et al. 2016). Physiological noise correction was either performed with Conn, which is a component based correction (Whitfield-Gabrieli & Nieto-Castanon 2012), or with Brainwaver, which makes use of a wavelet decomposition (Achard et al. 2006). A proportional threshold was applied on weighted (A) or binarized (B) connectivity matrices.

There were no group differences in age, gender, acquisition site or education (Table 1). A replication was also performed using an alternative subdivision strategy in which a selection of cohorts was made based on median age instead of pairwise matching for gender and age. This replication showed similar patterns, with Conn connectomes showing an AUC ranging from 0.56 to 0.70, compared to Brainwaver connectomes yielding 0.32 to 0.52 (Supplementary Figure 1). Overall functional connectivity (van den Heuvel *et al.*, 2017) was corrected by rescaling all elements of a matrix with respect to their contribution to overall functional connectivity on group level (Supplementary Figures 2-4). After application of this correction, the same patterns in the results were obtained for classification, with Conn connectomes yielding an AUC from 0.57 to 0.76 and Brainwaver connectomes 0.32 to 0.57 (Supplementary Figure 5). Exclusion of outliers in terms of mean framewise displacement did not have a considerable effect on the classification, with the AUC for Conn connectomes ranging from 0.56 to 0.73 compared to Brainwaver connectomes ranging from 0.30 to 0.51 (Supplementary Figures 6-7).

### Effect on modular organization

We then investigated how modular organization was affected by different connectome construction methods. The assignment of each node to a module was qualitatively compared and quantitative group differences in participation coefficient (PC) and within module-Z score were also investigated (see Materials and Methods).

Across all atlases, the nodes of modules for the Brainwaver connectomes showed less spatial proximity compared to Conn connectomes. For Brainwaver connectomes, nodes of the same module were more dispersed for both HD and healthy controls when compared to Conn connectomes (Figure 3). The Supplementary Materials include all lists of the module assignments for the three atlases (see also corresponding Supplementary Figures 9-11). The most important effect of physiological noise correction on modularity was observed in the functional atlas: for healthy controls, Brainwaver connectomes provided dispersed modules that were highly different in number and location across thresholds, while Conn connectomes showed nodes in spatial proximity that had a consistent modularity across thresholds (Figure 4). In addition, the functional atlas also gave rise to modules that were less demarcated due to its lower resolution (121 ROIs), compared to the structural and multi-modal atlases (316 and 360 ROIs, see Figure 5).

**Figure 3:**
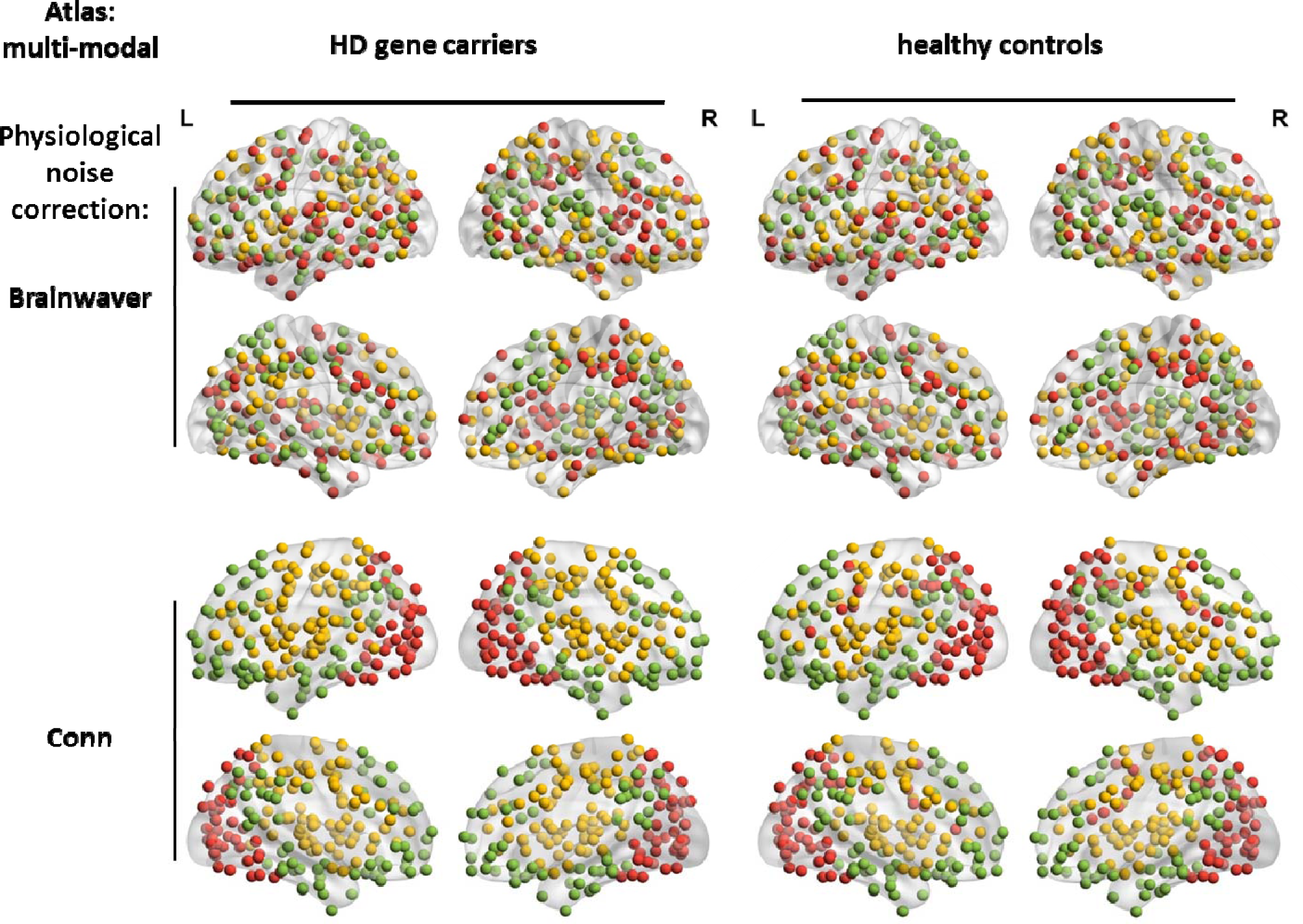
The nodes of the modules are more dispersed for Brainwaver connectomes, while anatomically coherent for Conn connectomes. The visualization of the group community affiliation vector for the HD gene carriers and healthy controls is based on weighted unthresholded matrices. In all instances, 3 modules were detected by the Louvain community detection algorithm. For Conn connectomes these modules seem to correspond with a posterior (red), fronto-temporal (green) and fronto-parietal network (yellow).

**Figure 4:**
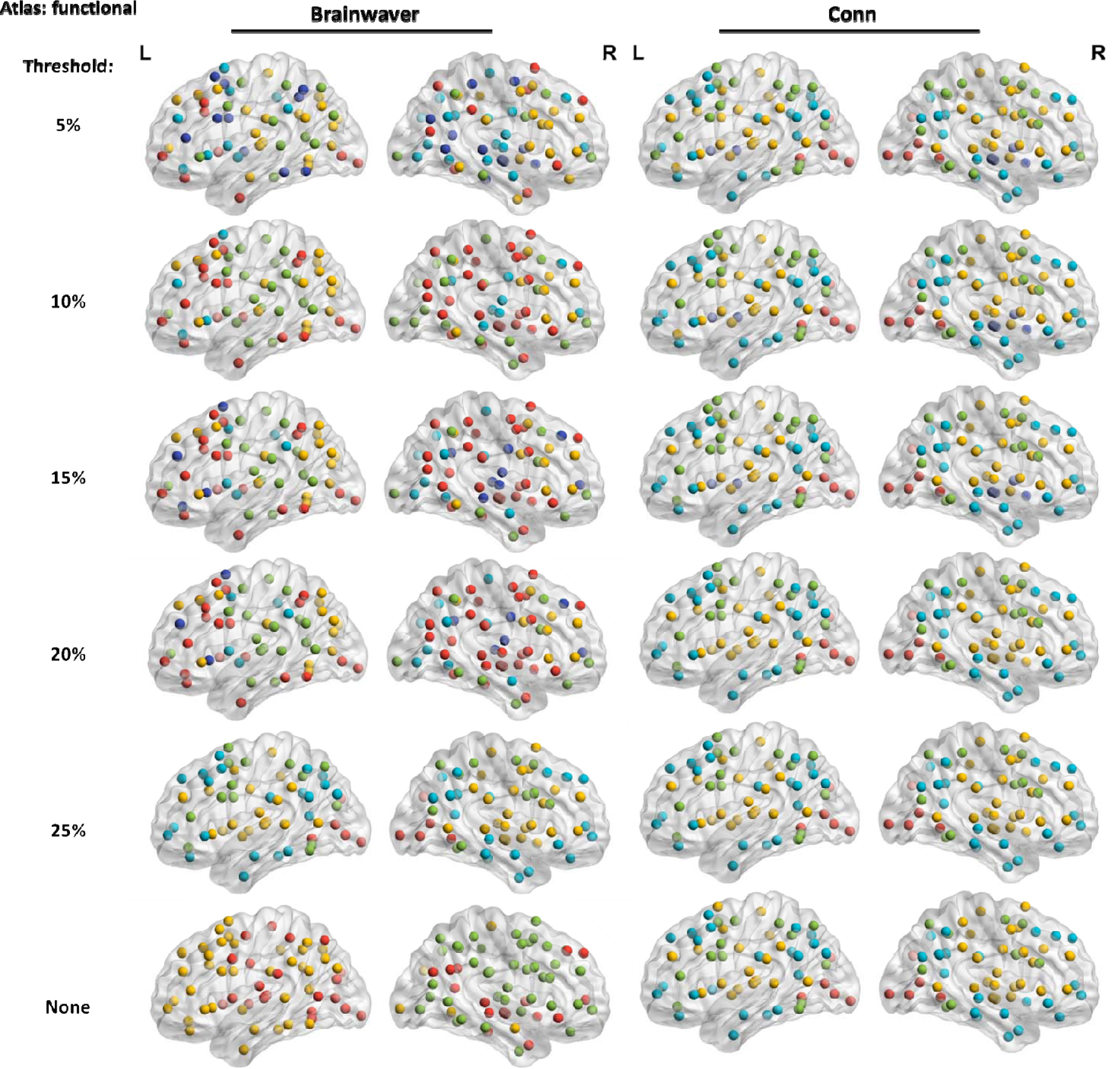
The modular organization is inconsistent in healthy controls across thresholds for Brainwaver connectomes, while consistent for Conn connectomes. The modules detected in weighted connectomes are visualized. A functional atlas was used for both types of physiological noise correction.

**Figure 5:**
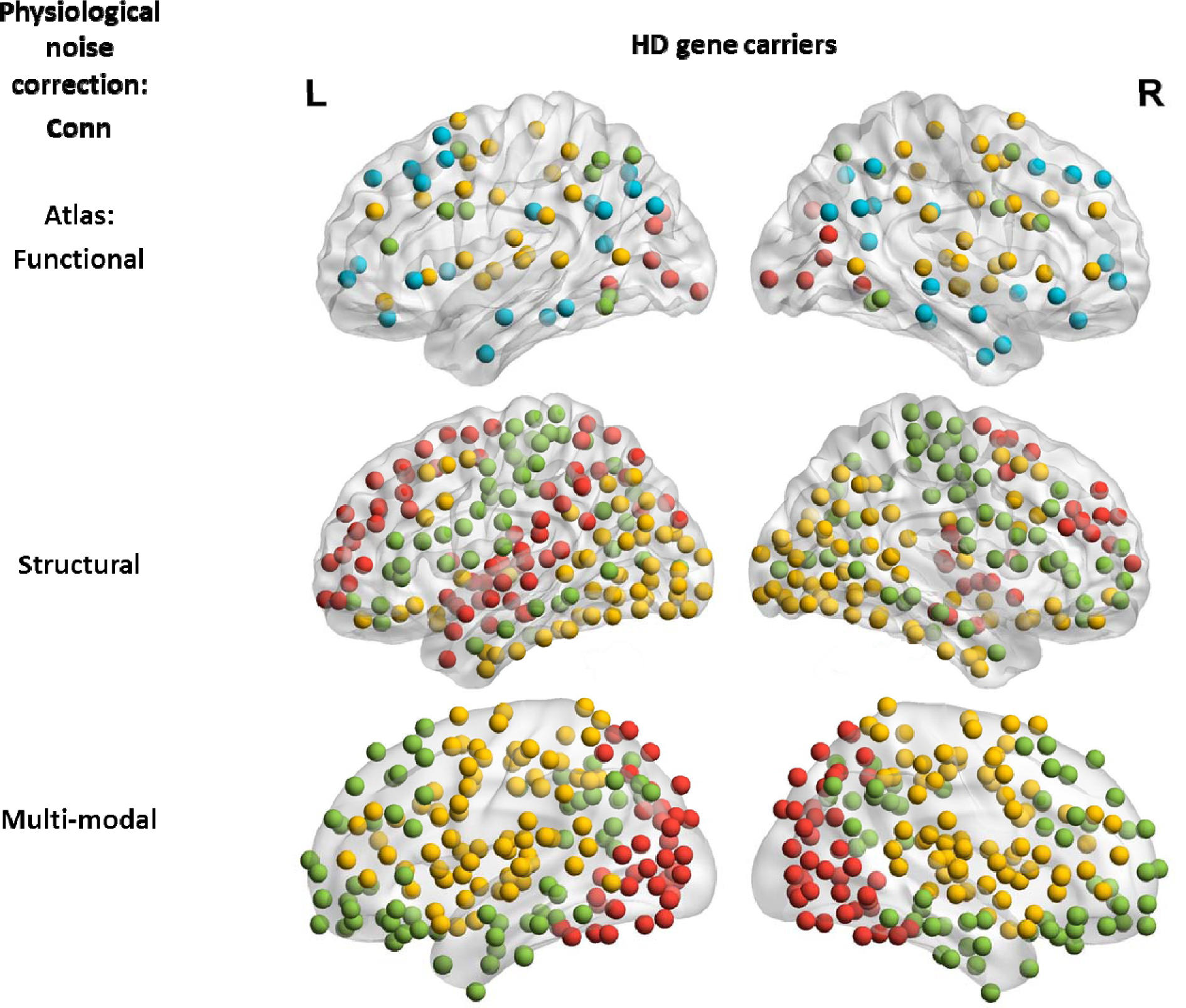
Comparison of modularity on HD gene carrier group level across atlases. The visualization of the group community affiliation vector for the HD gene carriers and healthy controls is based on weighted unthresholded matrices. There were 4 modules detected using the functional atlas (121 ROI), and 3 in the structural and multi-modal atlas (316 and 360 ROI). The distribution of the nodes of a module is less informative in a lower resolution atlas compared to higher resolution.

For quantitative assessment of modularity, nodes that were significantly different (p<0.05, FDR corrected) between HD and healthy controls were summed with respect to participation coefficient (PC) and within module-Z score. Table 2 shows that the type of physiological noise correction, the parcellation atlas, thresholding and binarization all had an important effect on modularity group differences.

**Table 2:**
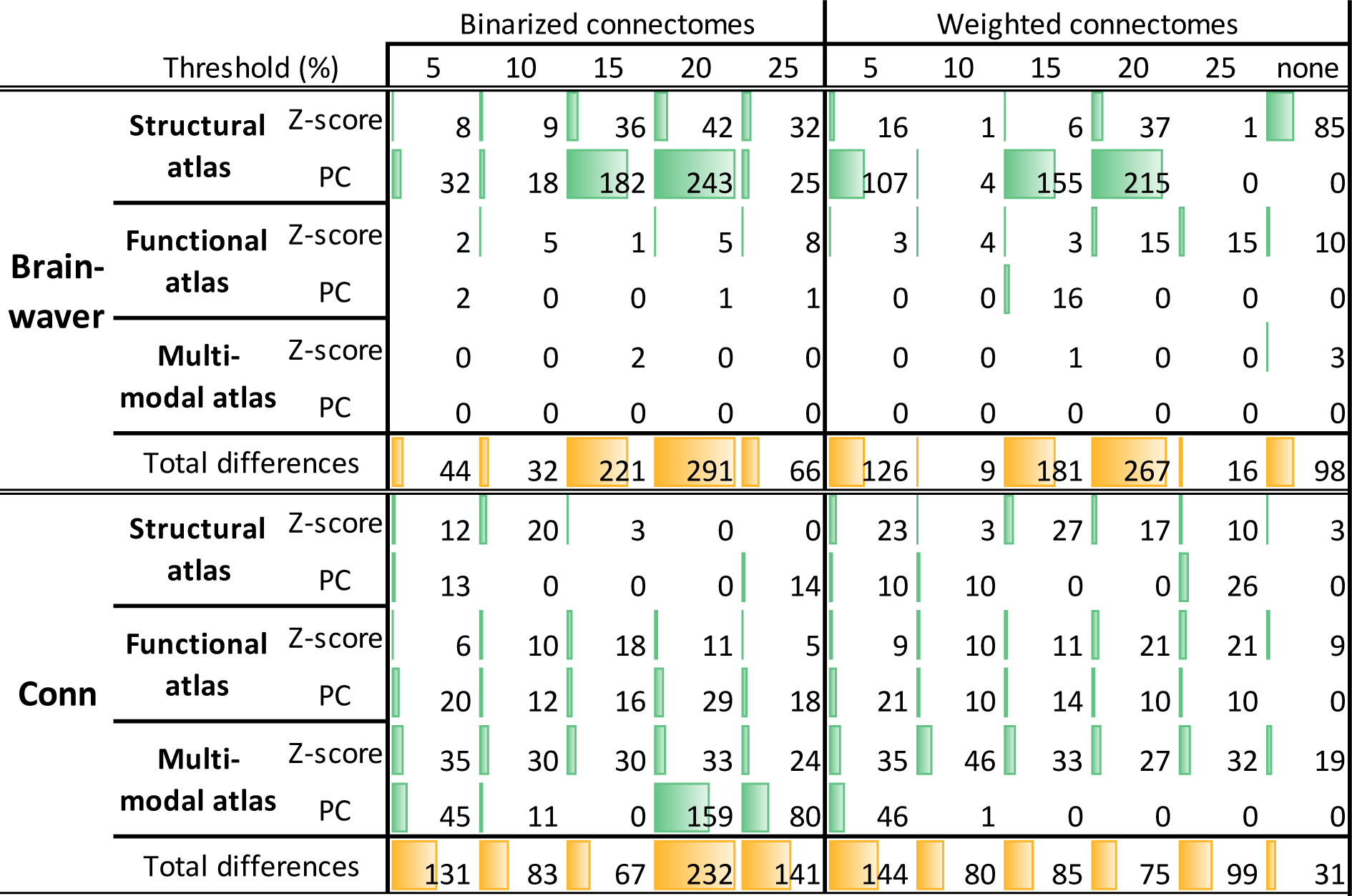
**Number of nodes with a different modularity property for Brainwaver and Conn connectomes.** For each connectome construction strategy, the connectomes were assessed for differences in participation coefficient (PC) or within module Z-score (Z-score) between HD gene carriers and healthy controls with permutation testing (10 000 iterations). The sums of all significant nodes (p-value < 0.05, FDR corrected) are reported, regardless of the direction of the effect.

First, the effect of thresholding and binarization was investigated for Conn connectomes using the multi-modal atlas. The weighted connectomes showed 46 significant nodes for PC at a threshold of 5% and only 1 node at threshold 10% and zero at thresholds of 15%, 20% or 25% (Figure 6a). After binarization an irregular pattern across the thresholds in both the number and location of significant nodes was observed, which showed that binarization had a substantial effect on group differences in modularity (Figure 6b).

**Figure 6:**
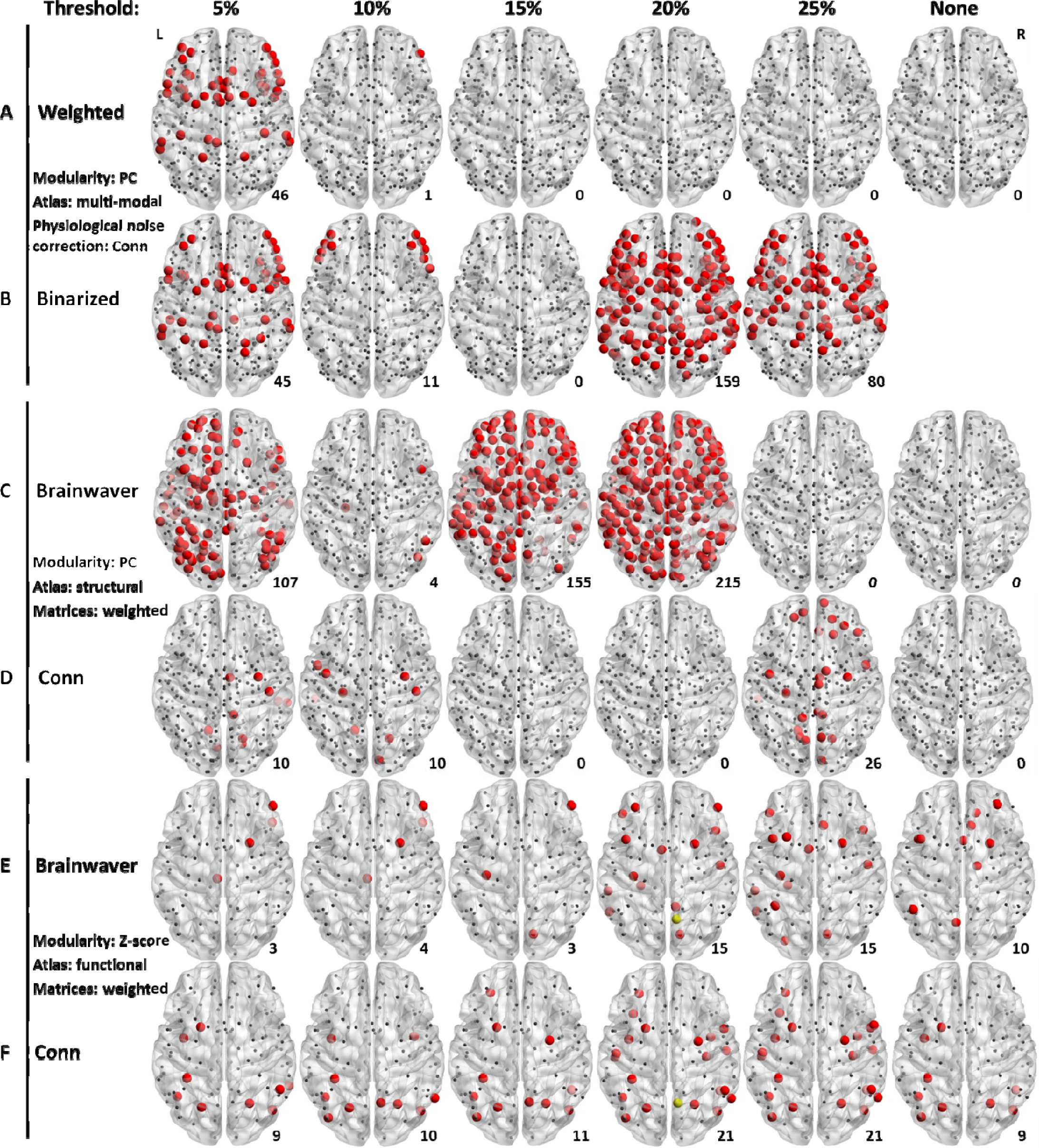
Number and location of nodes that differ modularity between HD and healthy controls are substantially effected by thresholding, binarization and type of physiological noise correction. The number of nodes that had differences in participation coefficient (PC) or module Z-score (Z-score) between HD gene carriers and healthy controls, are visualized in a dorsal view (permutation testing, p-value < 0.05, FDR corrected). This number is for each comparison given in the bottom right corner. Binarization and thresholding have a strong influence on detection of group difference: example shown for weighted connectivity matrices (A) and the binarized equivalent (B) using a multi-modal atlas. Connectomes after wavelet decomposition (Brainwaver) versus component-based correction (Conn) for physiological noise lead to substantially different outcomes (C and D): example shown for weighted matrices, using a structural atlas. When number of detected nodes are comparable, connectomes corrected with Brainwaver or Conn do not lead to overlap in location (E and F): example shown for a functional atlas, using weighted matrices. From all pairwise comparisons between E and F, only 1 single node is in common between two methods of physiological noise correction (yellow node, at threshold of 20%). For full results, see Table 2.

In addition to the influence of binarization and thresholding, the effect of physiological noise correction was also examined. For the structural atlas, Brainwaver connectomes showed the following number of PC significant differences: 107 nodes at 5%, 4 nodes at 10%, 155 nodes at 15%, 215 at 20% and no nodes were significant at a threshold of 25%, or when no threshold was applied. Conn showed 10 nodes at a threshold of 5% and 10%, zero nodes at 15% and 20%, 26 nodes at 25% and no nodes were found significant when no threshold was applied (Figure 6c-d).

In order to distinguish between the number of significant nodes and their location, the within module Z-score using the functional atlas was further investigated because both types of physiological noise correction yielded a similar number of nodes. Brainwaver showed 3 nodes at 5%, 4 nodes at 10%, 3 nodes at 15%, 15 nodes at 20% and 25%, and 10 nodes were found significant when no threshold was applied. Conn showed 9 nodes at 5%, 10 nodes at 10%, 11 nodes at 15%, 21 nodes at 20% and 25%, and 9 nodes when no threshold was applied. Regarding the location of significant nodes, there was no overlap when comparing Brainwaver and Conn connectomes. When no threshold was applied, the 10 nodes detected in the Brainwaver connectome did not overlap with the 9 nodes detected in the Conn connectome.

There was also no overlap in the nodes at the thresholds 5%, 10%, 15% or 25%. Only at a threshold of 20% one single node (showed in yellow) was in common from 15 and 21 nodes detected using Brainwaver and Conn connectomes respectively (Figure 6e-f).

## Discussion

The aim of this work was to perform a systematic comparison of connectome construction strategies to provide recommendations for representing functional networks in neurodegeneration. The following aspects of connectome construction strategies were investigated: principal component-based physiological noise correction versus wavelet decomposition, the type of parcellation atlas (functional, structural and multi-modal), weighted versus binarized networks, and unthresholded versus proportionally thresholded networks.

For the classification of HD gene carriers and healthy controls, the connectomes making use of principal component-based physiological noise correction yielded higher discrimination rates compared to a physiological noise correction with wavelet decomposition. When a functional atlas was used in combination with principal component-based correction, the highest discrimination rates were obtained. The operation of thresholding or binarization did not enhance classification performance. The classification was not driven by group differences in gender, age, education, site of acquisition, overall functional connectivity or motion. The findings were consistent across two strategies of cohort subdivision ensuring equal sample size.

The modular organization was most anatomically coherent and consistent across thresholds when using principal component-based physiological noise correction. Connectomes obtained after wavelet decomposition for physiological noise correction resulted in dispersed modules, and these modules were different again across thresholds when a functional atlas was used. While this could represent complete modular breakdown in HD, the observation of dispersed modules in healthy controls does not support this (Sporns, 2011), suggesting this is methodological rather than biological.

Group differences in modular organization in terms of participation coefficient and within module Z-score were highly affected by the type of physiological noise correction and by the threshold applied, whether or not combined with binarization. There was no relationship between the percentage of thresholding and the number of nodes that were significantly different between HD gene carriers and healthy controls. Moreover, steep changes in both the number of nodes and their location were observed when incrementally increasing the threshold.

Whereas thresholding and binarization did not increase the ability to discriminate between groups when classifying, these operations showed a strong effect on identification of modular group differences. This was not keeping in with former findings of the reliability of modular organization in thresholded matrices, be it with a larger sparsity (He *et al.*, 2009; Meunier, 2009), but was in agreement with the caution around thresholding (Scheinost *et al.*, 2012; Garrison *et al.*, 2015), and the relevance of weaker connections (Gallos, Makse and Sigman, 2012; Santarnecchi *et al.*, 2014; Goulas, Schaefer and Margulies, 2015). These findings together suggest that thresholding, whether or not combined with binarization, has a risk to introduce artificial group differences, especially when considering that there is no standard to choose the threshold. One way to overcome the arbitrariness of choosing a threshold and to enhance the reliable detection of network group differences, may be the use of unthresholded weighted matrices.

While machine learning can have high diagnostic accuracy in schizophrenia (Davatzikos *et al.*, 2005) or Alzheimer’s disease (Klöppel *et al.*, 2008), it is of limited utility because the diagnosis can usually be made clinically in such cases. However, if the diagnosis is known *a priori*, machine learning can be useful to indicate which imaging methodology yields the highest discrimination rate. In the present study, the diagnosis of Huntington’s disease was made with genetic testing so could be used as a gold standard, and so machine learning can indicate the most reliable connectome construction strategy. However, the robustness of the use of machine learning in connectomics is under further investigation (Brown and Hamarneh, 2016). The use of proposed null models for brain networks (Rubinov and Sporns, 2011; Hosseini and Kesler, 2013), test-retest analysis (Du *et al.*, 2015; Marchitelli *et al.*, 2016), exploring different confidence measures of classification prediction (Gammerman and Vovk, 2007; Nouretdinov *et al.*, 2011), different types of datasets and the use of different types of machine learning algorithms such as Scikit-learn or its recent neuroimaging equivalent Nilearn (Abraham *et al.*, 2014), are all but a few methods to assess the generality of application of machine learning in connectomics.

The assessment of modular organization is complimentary to machine learning classification, as it enables direct mapping of modules on the brain. For both of these methodologies, the functional atlas gave rise to the highest classification performances, and it was also the most stable atlas across thresholds in terms of modularity for Conn based connectomes. Potentially, this is due to the size of the atlas. This is important as currently there is no consensus on the most reliable type of parcellation atlas (Arslan *et al.*, 2017). Another explanation for the performance of the functional atlas could be that the multi-modal and structural atlas did not include sub-cortical regions, as for instance is the case for the multi-modal Brainnetome atlas (Fan *et al.*, 2016). However, the specific way how an atlas is composed, remains an issue for its application. For example, Arslan and colleagues cautioned that the Glasser multi-modal atlas may be positively biased toward the specific modalities (task fMRI, myelin content) used for its composition (Arslan *et al.*, 2017), and other atlases may suffer from the same biases. Recently, gradient-weighted Markov Random Field models have been used to account for these biases by detecting changes in functional connectivity similarities, showing that these multiresolution parcellations may be more homogenous than other multimodal atlases, including the Glasser atlas (Schaefer *et al.*, 2017).

One limitation of our study is that it does not provide a standard to assess which construction provides the best topological organization of the connectome. While replication of our findings in other neurodegenerative diseases will be necessary to generalize our current findings to a broader field, the strength of Huntington’s disease as a model for neurodegenerative diseases is that the diagnosis is technically robust and accurate. This implies that cohort of HD gene carriers consisted of both participants with manifest symptoms and participants at a pre-manifest stage, sometimes decades before the onset of disease. When assessing group differences in functional connectomes of HD gene carriers and controls, genetic testing thus served as a gold standard.

Our work suggests that principal component-based physiological noise correction outperformed wavelet decomposition. An in depth analysis of these and other types of physiological noise correction, such as Bayesian methods (Särkkä *et al.*, 2012) or independent component analysis (Griffanti *et al.*, 2014) is beyond the scope of this study. The novelty in this systematic comparison is that it combined the strengths of machine learning and investigation of modular organization. Our work also considered a variety of factors from different levels (type of parcellation atlas, physiological noise correction, choice of threshold and binarization) in connectome construction to assess their mutual effects on the connectome, instead of limiting the investigation to a single factor alone. A reproducibility analysis of our methodology would be welcome, to verify how consistent the results are across different datasets.

## Conclusions

We performed a systematic comparison of various connectome construction strategies based on resting state fMRI data from TrackOn-HD project. Principal component-based physiological noise correction resulted in a higher group discrimination rate, and the use of a functional atlas outperformed other atlases. We showed that the type of physiological noise correction, the parcellation atlas, thresholding and binarization all have a substantial effect on the detection of group differences based on modularity. This raises important methodological concerns for functional connectomics in general. The results of the present study support the use of unthresholded and weighted matrices in combination with principal component-based noise correction methods. While the use of a functional atlas may provide more consistent results, a meticulously made trade-off with higher resolution atlases may also be necessary.

## Acknowledgments

This work was supported by Wellcome Trust Grants (091593/Z/10/Z, 103437/Z/13/Z, to G.R. and P.MC and 200181/Z/15/Z to E.J., S.G., R.S. and S.T.) and the National Institute for Health Research, University College London Hospitals, Biomedical Research Centre. The Track-On HD study is funded by the CHDI Foundation, a not-for-profit organization dedicated to finding treatments for Huntington’s disease.

## Track-On HD investigators

The Track-On HD Investigators are A. Coleman, J. Decolongon, M. Fan, T. Petkau (University of British Columbia, Vancouver, Canada); C. Jauffret,D. Justo, S. Lehericy, K. Nigaud, R. Valabrègue (ICM and APHP, Pitié-Salpêtrière University Hospital, Paris, France). A. Schoonderbeek,E. P. ‘t Hart (Leiden University Medical Centre, Leiden, the Netherlands);D. J. Hensman Moss, R. Ghosh, H. Crawford, M. Papoutsi, C. Berna,D. Mahaleskshmi (University College London, London, United Kingdom). R. Reilmann, N. Weber (George Huntington Institute, Münster, Germany); I. Labuschagne, J. Stout (Monash University, Melbourne, Australia); B. Landwehrmeyer, M. Orth, I. Mayer (University of Ulm, Ulm, Germany); H. Johnson (University of Iowa, Iowa City, IA); D. Crawfurd (University of Manchester, Manchester, United Kingdom).

## Abbreviations

AUC: area under the curve
BCT: Brain Connectivity Toolbox
BW: Brainwaver
CM: connectivity matrix/matrices
CompCor: principal component-based noise correction
CSF: cerebrospinal fluid
FC: functional connectivity
FDR: false discovery rate
HC: healthy controls
HD: Huntington’s disease
MODWT: maximal overlap discrete wavelet transform
MST: minimum spanning tree
PC: participation coefficient
PRoNTo: Pattern Recognition for Neuroimaging Toolbox
ROI: region of interest
rs-fMRI: resting state functional Magnetic Resonance Imaging
SD: standard deviation
SPM: statistical parametric mapping
T1WI: T1-weighted imaging
WMW: Wilcoxon-Mann-Whitney test

